# Can blood at adult age predict epigenetic changes of the brain during fetal stages?

**DOI:** 10.1101/2022.11.28.518197

**Authors:** Monica Strawn, Timothy J. Safranski, Susanta K Behura

**Affiliations:** Division of Animal Sciences, University of Missouri, Columbia Missouri 65211; MU Institute for Data Science and Informatics, University of Missouri, Columbia Missouri 65211; Interdisciplinary Neuroscience Program, University of Missouri, Columbia Missouri 65211

**Keywords:** fetal brain, sex, pig, methylation, blood, adult

## Abstract

Correspondence in DNA methylation between blood and brain is known in humans. If this pattern is present in pig has not been examined. In this study, we profiled DNA methylation of blood from pigs at adult ages, and compared those with the methylation profiles of fetal brain. Neural network regression modeling showed specific methylations in the adult blood that can reliably predict methylation of the fetal brain. Genes associated with these predictive methylations included markers of specific cell types of blood and brain, in particular, markers of bone marrow hematopoietic progenitors, and glial cells primarily the ependymal and Schwann cells of brain. The results of this study show that developmental methylation changes of the brain during fetal stages are maintained as an epigenetic memory in the blood in adult life. Thus, pig models may be harnessed to uncover potential roles of epigenetic memory in brain health and diseases.

## Background

The pig (*Sus scrofa*) is considered as a better animal model than rodents in studying human brain [1] and is increasingly used to study developmental disorders and neurodegenerative diseases of brain [2–6]. Imaging studies show remarkable similarities in development between pig and human fetal brain [7]. Pig brain is gyrencephalic similar to that of human [8], and shows similar patterns of neuronal networks and subcortical nuclei seen in nonhuman primates and humans [9]. It has similar white to gray matter ratio like human [10].

Our recent study identified genes differentially regulated (data accession # GSE178970) during development of fetal brain in pig [11]. That study further found common genes that were expressed in both human and pig developing brain in support of earlier report [12]. Studies by other have shown that gene expression in pig brain is influenced by various epigenetic factors including chromatin modification and DNA methylation [13]. Recently, we studied epigenetic changes during development of fetal brain in pig [14]. Multi-tissue analysis by others has identified methylation sites associated with aging in pig [15]. Studies have also shown signatures of similar changes in blood DNA methylation during development and aging [15–18] suggesting their role in epigenetic clocks of individual organs [19]. Furthermore, studies have shown that there is correspondence in DNA methylation between brain and blood in human [20], and specific methylations of blood can be reliable biomarkers of neurodegeneration and neuropsychiatric diseases [21–23]. However, no study has been performed to test if DNA methylation of blood at adult ages can inform methylation changes that occurred in the brain at the fetal stages. The aim of the current study is to test the hypothesis that epigenetic memory [24–27] of brain development in the fetal life persists in the blood DNA in adult life.

## Results

### Methylation changes of blood DNA at adult ages

Whole-genome methylation profiling was performed of blood collected from pigs of both sexes at three adult ages, months (mo) 10, 12 and 21. Data analysis showed increasing number of CpG methylation with increase of adult age (**Figure 1**). However, a common core of 96,361 methylation sites was persistently observed in the blood of both sexes at the three adult ages indicating that these methylations are persistent in adult blood (henceforth referred to as Adult Blood Methylations or ABMs). Though ABMs were associated the common CpG sites in both sexes at the three adult ages, level of methylation of those sites varied between ages and sexes (**Figure 1**).

**Figure 1.**
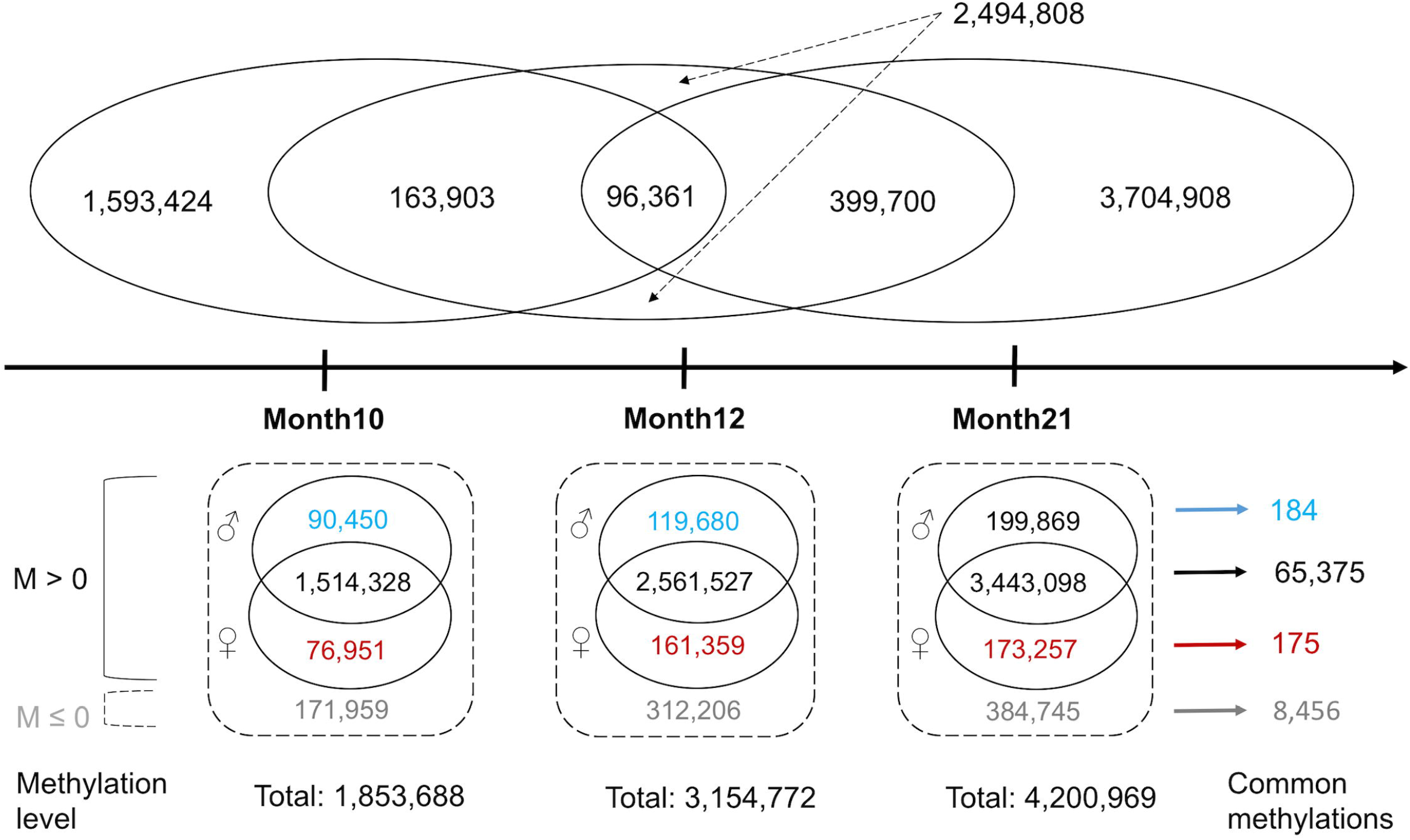
Changes in methylation of blood at adult ages. The upper panel shows the number of methylation sites that are either specific or common to the three adult ages (in months). The lower panel shows methylation level (measured by M-values) between male vs. female adults. The numbers shown in blue represent sites with M > 0 in male blood but M ≤ 0 in female blood. The numbers shown in red represent sites with M > 0 in female blood but M ≤ 0 in male blood. The numbers shown in black (common) represent M > 0 sites in both sexes. The numbers shown in grey outside the circles represent methylation sites with M ≤ 0 in both sexes. The number of common methylation sites in each of these four groups across the three ages are shown right to the arrows.

### Fetal brain and adult blood have common methylations

We compared blood methylations of adult blood with methylation profiles from developing fetal brain at three gestation days (GDs) 46, 60 and 90 generated from our earlier study (accession number GSE178983) [14]. The fetal brain methylation data showed a common core of 69,757 methylation sites was observed across the three GDs in both sexes (**Supplementary Figure 1**) indicating that these methylations were persistent during fetal brain development (henceforth referred to as Fetal Brain Methylations or FBMs).

Comparison between ABMs and FBMs showed that 42,250 methylation sites were common between ABMs and FBMs (**Figure 2A**). These common sites were methylated in the brain at fetal stages and blood of at adult stages in both sexes, henceforth referred to as Blood Brain Methylations (BBMs) that are listed in **Supplementary Table 1**. BBMs showed variable chromosomal distribution in the pig genome (**Figure 2B**). In particular, specific regions of Chr4, Chr10, Chr14 and Chr17 showed higher incidences of BBMs (> 3,000 per 30Mb DNA) than other parts of the genome. Many BBMs were localized within CpG islands (n=1,784) and CpG shores (4,796) (**Supplementary Table 2**). We observed variable lengths of physical clusters of methylation sites from shore to island (‘methylation bridges’) in specific chromosomal regions of Chr7, Chr10, Chr12 and Chr14 listed in **Supplementary Table 2**. The methylation bridges in Chr10 (position 14001133 to 14007361) and Chr12 (from position 58563865 to 58572915) were localized within long non-coding RNAs (lncRNA), though DNA methylation of noncoding RNA genes is not uncommon [28].

**Figure 2.**
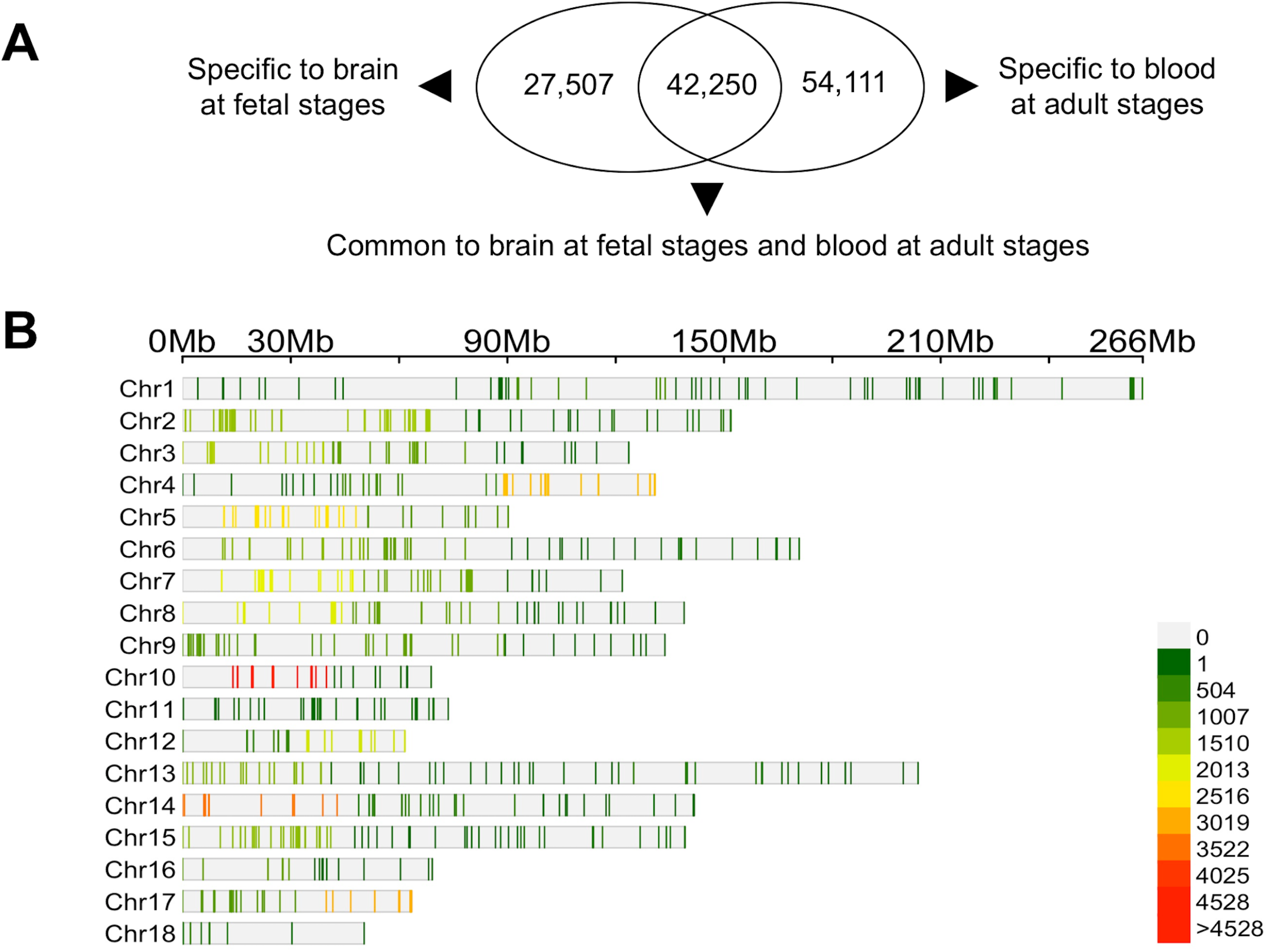
Comparison of methylation sites between fetal brain and adult blood. **A**. Venn diagram shows number of methylation sites either specific or common to the fetal brain and adult blood. Note that this comparison was made between the 69,757 common methylation sites in the fetal brain and 96,361 common methylation sites in the adult blood. **B**. Per-chromosome density map of methylation sites (n=42,250). These are the sites that are methylated commonly in fetal brain and adult blood. The 30Mb bins per each chromosome is shown on the top. The color in the density map corresponds to the scale showing color codes for different number of methylation sites.

### Functional annotation of genes methylated in both fetal brain and adult blood

We identified a total of 199 genes where BBMs were found either in the gene body or immediate (within 1 kb) flanking DNA (**Supplementary Table 3**). Several of these (n=29) genes were related to ‘transport’ function suggesting the relevance of transport genes in brain development and physiology [29]. These genes also included several known markers of blood as well as brain cells. In particular, these genes were markers of bone marrow hematopoietic progenitors and glial cells of the brain, primarily the Ependymal and Schwann cells (**Supplementary Table 3)**.

### Canonical correlation in DNA methylation between glial cells of brain and progenitor cells of adult blood

We identified specific methylations (n=1,456), including those associated with the marker genes of blood progenitor and brain glial cells by bicluster analysis [30] where methylation level varied in four distinct clusters (**Figure 3**). Bicluster analysis identifies clusters taking both rows and columns (in our case, rows are methylations, and columns are the blood and brain samples) into consideration unlike the conventional cluster analysis that considers changes either among the rows or columns. Besides the four biclusters, we identified another cluster of methylations (n=102), but it didn’t show bicluster pattern among all the blood and brain samples. Thus, it was excluded from further analysis. The covariates (product of correlation and variance) of the methylation changes, based on canonical correlation analysis [31], for the four biclusters are listed in **Supplementary Table 4**. Pair-wise comparison was performed based on cluster distances of methylation changes between the bicluster groups along with the non-cluster methylations (**Supplementary Figure 2**). It showed that the methylations among the bicluster groups were highly correlated (Pearson r > 0.96) among each other. The non-cluster methylations showed an average correlation of 0.63. Principle component analysis further showed distinct grouping patterns of the brain and blood samples between bicluster and non-bicluster methylations (**Supplementary Figure 3**).

**Figure 3.**
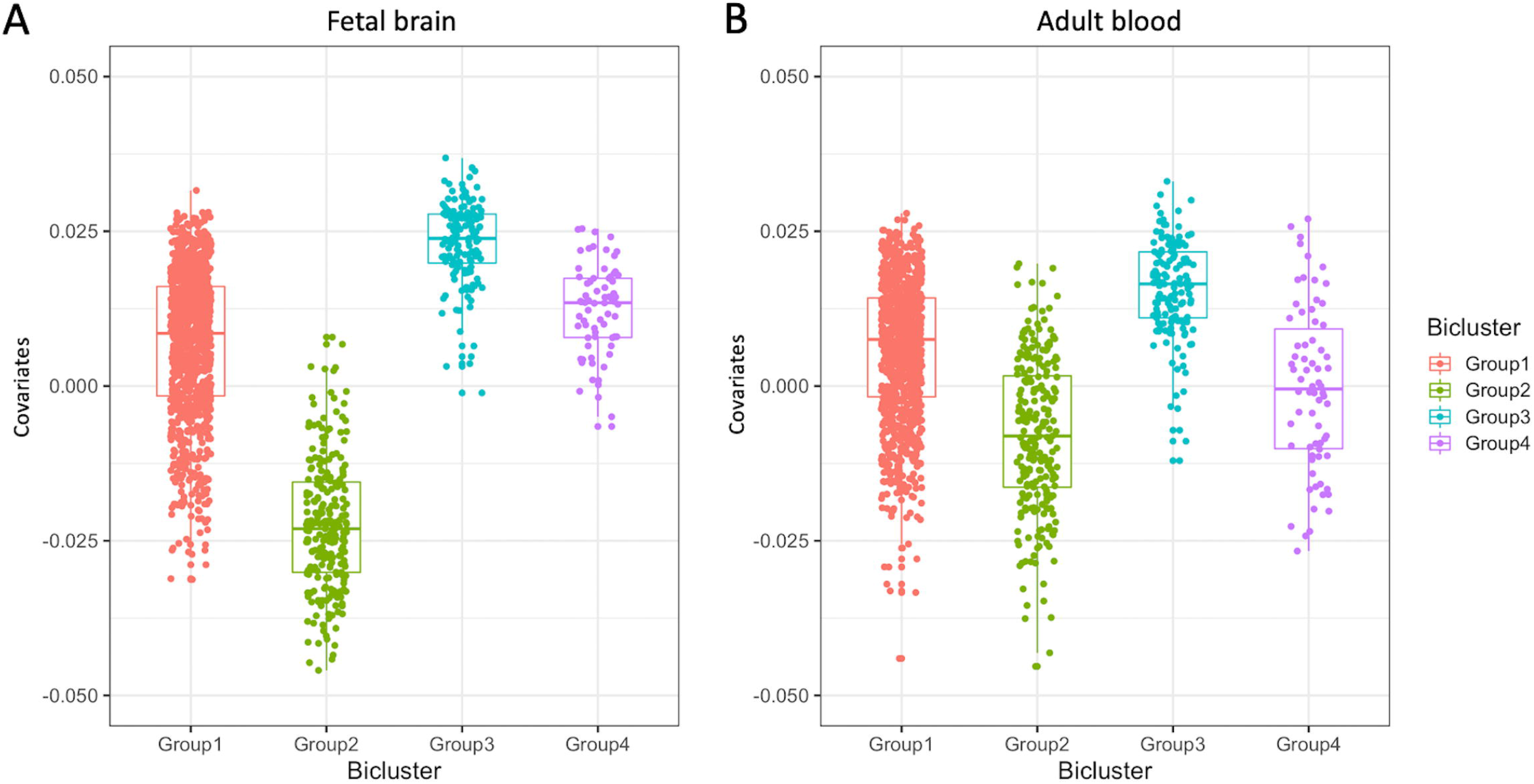
Boxplot showing four groups (clusters) of CpG sites that are methylated in a canonical correlation manner between fetal brain and adult blood samples. The x-axis shows the groups and y-axis shows the major covariate of methylation level of the sites in the fetal brain (**A**) and adult blood (**B**). The horizontal line within each group (color coded) represents mean value of covariates for each group.

We applied neural network models, as described in our earlier study [32], that learnt methylation changes in adult blood and predicted methylation changes of brain during fetal development. First, we applied neural network for data classification of methylation in fetal brain and adult blood. The classification was based on whether the methylations were biclustered (assigned 1 as predicted value), or not (assigned 0) between fetal brain and adult blood (**Supplementary Figure 4**). In these analyses, 10% of the methylation sites were randomly used as training data to develop the model, and then the trained models was used to compute the predicted variable. The predicted and observed values were then compared. These within-group analysis showed more than 95% accuracy in classifying methylation data both in fetal brain as well as adult brain. Then, we applied neural network regression analysis to predict changes in methylation changes in fetal brain based on changes in the adult blood. In this analysis, the model learnt the methylation changes in the aging adult blood as predictor variables relative to the canonical covariates as predicted variables. The trained model was then used to predict canonical covariates of fetal brain based on methylation changes at different developmental stages. The predicted values showed a significant positive correlation (Pearson r = 0.961) with the observed values in the fetal brain suggesting that methylations in the blood DNA in the adult stage can reliably predict how methylation changed in the brain during fetal stages. Random sampling of the canonically correlated methylation sites was analyzed to assess mutual information [33] in fetal brain and adult blood separately. The network plots (**Figure 4**, also see **Supplementary Figure 5**) showed that methylation of these sites changes in a coordinated manners in the brain during fetal development and then in blood in adult ages.

**Figure 4.**
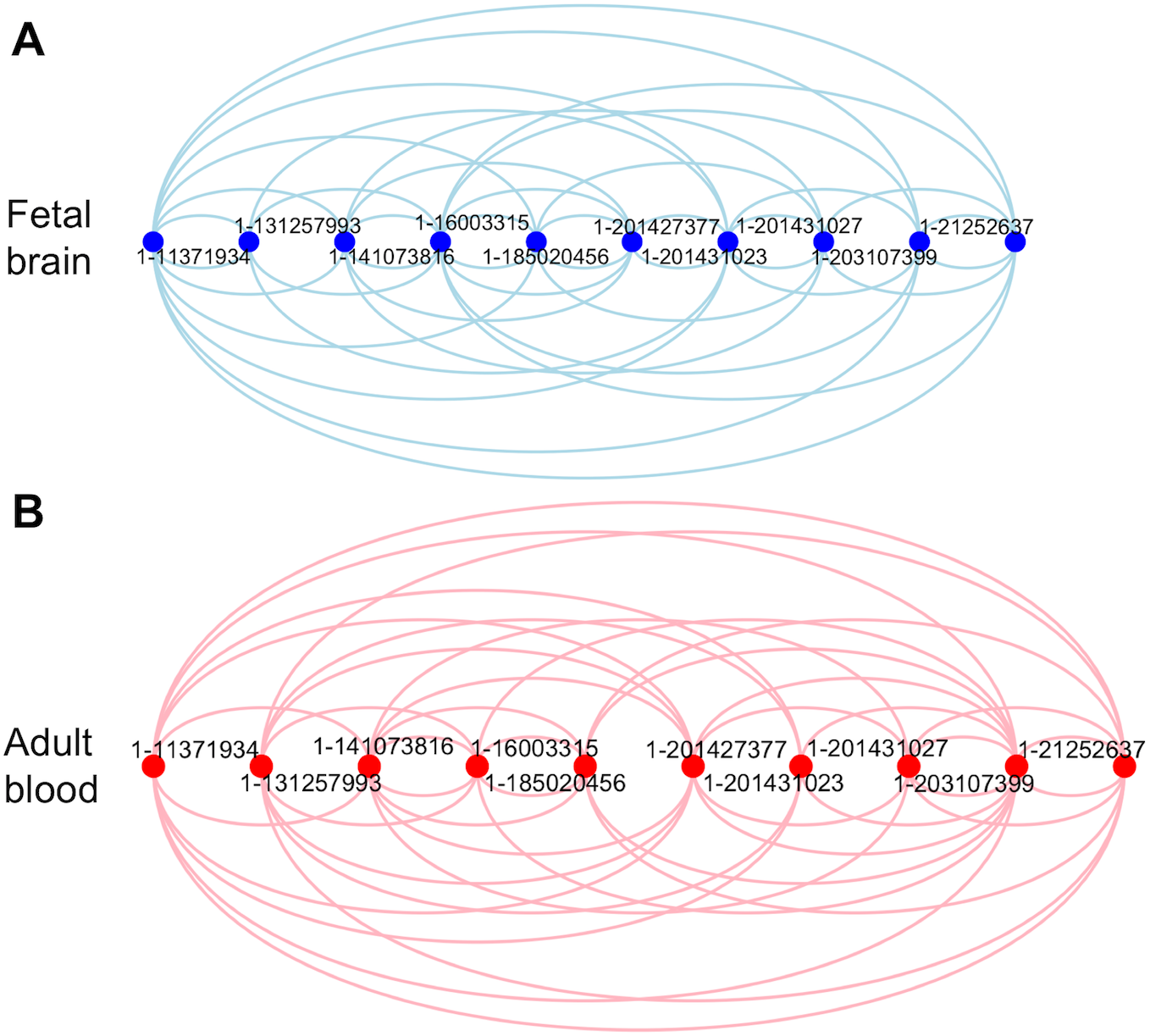
Arch plot showing pair-wise mutual information network among common methylation sites (n=10, shown as dot with labels representing the chromosome number followed by position of CpG site) in fetal brain (**A**) and adult blood (**B**). Each arch connecting two methylation sites represents the mutual information that is shared between variation of methylation at the two sites.

## Discussion

Emerging evidence suggest that epigenetic memory is a robust regulatory process that control various biological process including developmental cues for cell differentiation and lineage specification, adaptive cues to trigger response to intrinsic or environmental stress and inheritance cues to pass on parental exposures and experiences to the offspring [24,27]. In particular, role of epigenetic memory in hematopoietic stem cells in controlling diversity of cellular behavior as well as adaptation to inflammatory injuries are well established [25,26]. But if developmental events in the fetal life are embedded as epigenetic memory to persist in the blood in adult life remains unknown, which was the primary objective of the present study.

By comparing genome-wide methylation changes of blood in adult stages relative to methylation changes of the brain during fetal stages, we identified methylations in the adult blood that were predictive of methylations of fetal brain. Functional annotation of the genes associated with those methylations identified marker genes of blood progenitor cells and different glial cells of the brain. We identified common methylations in genes such as *KIT, PRRC2C, PRC1*, and *SEC61A* in both fetal brain as well as blood DNA. These genes are known markers of bone marrow hematopoietic progenitors, B lymphocytes, and lymphoid progenitor cells in blood, and the same genes are also markers of specific glial cells of the brain [34]. These findings suggested that DNA methylation differences in adult blood have significant concordance with fetal brain, a finding that is consistent with the idea of blood cells as keepers of epigenetic memory of early-life development [35,36]. We also used gene expression data of developing fetal brain of pig from our earlier study [11] to understand effect of the common methylations on gene regulation of brain during fetal development. The results of this analysis suggested these methylations occurred largely in genes that significantly altered prior to the rapid growth of fetal brain which occurred after gestation day 60. We have recently shown the importance of DNA methylation in gene regulation of fetal brain development in pig [14]. In addition, several genes were also related to transport function. Transport of ions and solute carriers are critical players that influence brain physiology and function [37]. In particular, solute carrier transporters in blood play major functions in the regulation of the blood-brain barrier [38] suggesting that methylation of transporter genes is a key epigenetic mechanism that persists both in brain at fetal stages and blood at adult ages. Several earlier studies have shown correlation in methylation changes between unrelated tissues including blood and brain [18,39–49]. Literature evidence suggests that methylation levels are generally highly correlated (r = 0.90) between blood and brain [45]. Earlier studies have shown that epigenetics play pivotal roles in fetal programming of adult heath [50]. Specific methylations are introduced in the genome at early developmental stages that persist through adult ages [51,52]. Evidence to support epigenetic programming of fetal brain development is known [53].

In general, methylation occurs in a tissue-specific manner [54]. However, concordance in methylation between blood and brain during aging is well known [17]. Edgar et al. (2017) curated the methylation sites in blood that correlates with methylation in brain (‘BECon’ or Blood–Brain Epigenetic Concordance tool) [55]. However, a key finding of the present study is the canonical correlation in methylation between fetal brain and adult blood which supports the idea that blood DNA methylation can be used as biomarkers of early development [54]. Moreover, methylation in blood occurs in age correlated manner [15,17,18,56]. Consistent with these earlier studies, the findings of the current study suggest that epigenetic changes in the blood DNA at adult ages can be used as predictors of epigenetic changes in the fetal brain. Further research is however needed to determine if blood stem cells in the adult contain the epigenetic memory of brain cells in the fetal stages due to adverse stress conditions during gestation. Maternal stress is known to cause different epigenetic disorders of the developing fetal brain [57–60]. Diagnosis of epigenetic disorders can be risky to fetal survival when the tests are performed with fetal samples directly. However, it is presently not known if methylation changes in blood are correlated with that of fetal brain due to pregnancy-related stresses. Thus, addition work are required to establish that possibility that can be a major step in advancing ongoing efforts in non-invasive prenatal screening of epigenetic disorders [61–64].

## Materials and methods

### Pigs and blood collection

The pigs used in this study were obtained from the University of Missouri Swine Research Complex (Columbia, MO). Blood was collected from age-matched boars and sows. At the time of collection, the boar and sow pairs were 10 months or 12 months or 21 months old. Blood was collected from three animals of either sex for each time point. The animal was restrained using a snare and whole blood was collected with a syringe and needle from the jugular vein and put in ethylenediaminetetraacetic acid (EDTA) tubes. Blood tubes were kept on ice and transported to the lab for DNA extraction. All research was conducted according to the Animal Care and Use Protocol approved by the University of Missouri Animal Care and Use Committee (ACUC).

### DNA extraction from blood

DNA from blood was isolated using QIAamp DNA Blood Mini Kit (Qiagen, Cat No./ID: 51104) as per manufacturer’s instruction. A total volume of 60μl DNA was eluted from each sample using spun columns provided in the kit. Concentration and purity of DNA was determined using Qubit 2.0 Fluorometer (Life Technologies), and an aliquot of DNA was also run in a 1.2 ethidium bromide stained agarose gel and imaged on ChemiDoc™ Touch Imaging System (Bio-Rad). Each sample was diluted to 100ng/μl using nuclease-free water and equimolar amount of DNA (300ng in 3 ul each) to generate 6 pooled DNA of adult blood that represented the three adult ages of both sexes. The advantages of using of pooled DNA in genome-wide methylation analysis has been described elsewhere [65].

### Genome-wide methylation analysis

Genome-wide methylation profiling of the blood samples was performed by enzymatic methyl-seq (EM-seq) using NEBNext^®^ Ultra™ II DNA Library Prep Kit. The use of pooled DNA generates accurate estimates of average level of methylation of each group, a detailed description of which is published earlier [65]. We used EM-seq approach that has many advantages over bisulfite conversion of DNA in whole genome bisulfite sequencing (WGBS) including superior sensitivity of detection of methylation, reduced DNA damage, low (pictograms) input DNA, uniform GC coverage and others [66–68]. Library preparation and sequencing were performed at the University of Missouri DNA Core Facility. The libraries were sequenced to 20x genome coverage using NovaSeq 6000.

### Bioinformatics analysis of EM-seq data

The raw sequences were assessed for quality with FastQC. TrimGalore was used to remove adapters and perform base quality trimming. The reads obtained from the quality control steps were mapped to the reference genome of swine, Sscrofa11.1, using Bismark [69]. After alignment, the methylation sites were extracted by using the *bismark_methylation_extractor* algorithm implemented within *Bismark,* and to generate the coverage files for methylation sites. As DNA methylation in mammalian somatic tissues mostly occur in cytosine-guanine (CpG) sites [70], only the CpG methylations were considered in this study. The coverage files contained information for methylation sites including the chromosome, start position, end position, methylation percentage, count methylated, and count unmethylated data. They were first processed to remove sites that were persistently low in mapped read counts (< 8) across samples, a filtration criterion generally used in methylation data analysis [71]. The of methylated and unmethylated reads counts were further filtered to remove sites that were either always methylated across samples or never methylated in any sample. In addition, methylations in chromosome X and Y were excluded throughout this study as they could confound bias in data analysis [72,73]. The methylated and unmethylated reads within a library were treated as a unit. Hence library size of each sample was considered as the average of the total number of methylated and unmethylated reads. Differential methylation analysis was then performed by fitting the normalized count data to generalized linear model using the *glmFit* function implemented in edgeR [71]. A likelihood ratio test was conducted using the *glmLRT* function to identify the differential methylation sites. The *topTags* function was used to extract the significant methylation sites with False Discovery Rate (FDR) < 0.05. All genomic arithmetic was performed using functions implemented in BEDTools [74].

### Statistical analysis

All statistical analyses were performed using *R*. The comparative analyses of the whole-genome methylation data of adult blood and fetal from a total of 12 animals that included 6 piglets and 6 adult pigs. The methylation level of each sites was calculated as M-values for all comparative analysis as these measures are less associated with heteroscedasticity for comparative analysis [75]. The M-value threshold equal to 0 represents that 50% of reads are methylated and the other 50% are unmethylated for each cytosine. We used this threshold to make an unbiased comparison of methylation sites between the fetal brain and adult blood. This is because we observed a greater number of sites with M > 0 in the adult blood than fetal brain. Thus, using a methylation greater than 50% would bias the analysis to adult blood. Similarly, using a methylation less than 50% would bias the analysis to fetal brain. Thus, we choose 50% as the methylation threshold for comparison of methylations between blood and brain. FPKM (fragment per kilo base per million) values [76] were used as measures of gene expression data (GSE178970). The variation in M-values and FPKM-values were measured by Euclidean distance. The distance matrix was used to perform hierarchical cluster analysis using *hclust* base function in R. The canonical correlation of M-values between fetal brain and adult blood samples was analyzed using *cancor* function in R package CCA.

### Predictive modeling

To explore if we can use adult blood methylation data to predict methylation changes that occurred in the brain during fetal development, we applied a predictive modeling approach using neural network analysis [77]. The R package *neuralnet* was used to perform the analysis. Data classification of methylation in fetal brain and adult blood was based on whether the methylated site showed bicluster pattern (assigned 1 as predicted value), or not (assigned 0). For training data, 10% of the methylation sites were randomly used. After the trained models compute the predicted variable, the predicted and observed values were compared. Neural network regression analysis was applied to learnt methylation changes in the blood during adult ages. In this analysis, the canonical covariates was used as predicted variables. The trained model was then used to compute predicted variable based on methylation changes during fetal developmental stages followed by correlation analysis of the predicted and observed values. The mutual information network analysis was performed using *minet* [78] using mutual information [33] of variation of methylation among the CpG sites in fetal brain and adult blood.

### Marker analysis of blood and brain cell types

Genes identified from the elastic net modeling were compared with dataset of known marker genes of brain and blood cell types. The dataset of marker genes of blood and brain cells were obtained from published single-cell transcriptomics studies [34,79–87]. The marker gene compression was performed as described in our earlier study [11].

## Acknowledgements

The authors are thankful to Jason L. Dowell for assistance in breeding the pigs used in this study. Library preparation and sequencing was performed at University of Missouri DNA Core. The authors acknowledge excellent help from Nathan J. Bivens and Mingyi Zhou of the DNA Core.

## Availability of data and material and Accession numbers

All the raw and processed data of this study have been submitted to the Gene Expression

Omnibus database under the accession number GSE178983.

## Competing interests

The authors declare that they have no competing interests

## Funding

This research was supported in parts by funding from the start-up grant from the University of Missouri, Columbia (SKB).

## Authors’ contributions

SKB designed the study, MS performed experiments, MS and SKB performed data analysis, MS,TJS and SKB wrote the paper.

## Supplementary Figures and Tables

**Supplementary Figure 1**. Methylation sites common in the fetal brain and adult blood that are localized either within gene body or in the immediate flanking DNA (within 1 kb) of genes. The genes related to transport function are indicated.

**Supplementary Figure 2.** Pair-wise correlation plot of methylation changes in fetal brain and adult blood for sites belong to the four biclusters (Groups) and sites that failed to cluster (Non-cluster). The scale on the right shows color coded bar for level of correlation.

**Supplementary Figure 3.** Comparison of cluster (**A**) and non-cluster (**B**) methylations in the fetal brain and adult blood samples by principal component analysis.

**Supplementary Figure 4.** Neural network plots of A) methylation of developing fetal brain (d=day) relative to bicluster relationship with adult blood, and B) methylation of blood DNA at adult ages (m=month) relative to bicluster relationship with fetal brain. The mean error and number of steps of the respective models are also shown.

**Supplementary Figure 5.** Arch plot showing pair-wise mutual information network among common methylation sites (n=10, shown as dot with labels representing the chromosome number followed by position of CpG site) in fetal brain (blue) and adult blood (pink) for different genomic regions (A, B, C). Each arch connecting two methylation sites represents the mutual information that is shared between variation of methylation at the two sites.

**Supplementary Table 1.** Changes in methylation of fetal brain. The upper panel shows the number of methylation sites that are either specific or common to the three gestation days. The lower panel shows methylation level (measured by M-values) between male vs. female fetuses. The numbers shown in blue represent sites with M > 0 in male fetal brain but M ≤ 0 in female brain. The numbers shown in red represent sites with M > 0 in female fetal brain but M ≤ 0 in male brain. The numbers shown in black (common) represent M > 0 sites in both sexes. The numbers shown in grey outside the circles represent methylation sites with M ≤ 0 in both sexes. The number of common methylation sites in each of these four groups across the three GDs are shown right to the arrows.

**Supplementary Table 2**. List of methylation sites common between fetal brain and adult blood. The chromosome number and position are listed for each methylation. Also, M-values are provided for each site.

**Supplementary Table 3**. Number of common methylation sites between fetal brain and adult blood found in CpG islands, shores, and bridges.

**Supplementary Table 4**. List of CpG sites (n=1,456) where methylation varied in a canonical correlation manner between fetal brain and adult blood in four groups (Groups 1 through 4). The estimates of covariates of methylation changes in fetal brain and adult blood are shown.

## Notes

### Competing Interest Statement

The authors have declared no competing interest.

