## Supplementary figures and images for "Can blood at adult age predict epigenetic changes of the brain during fetal stages?"

### Supplementary Figure 1

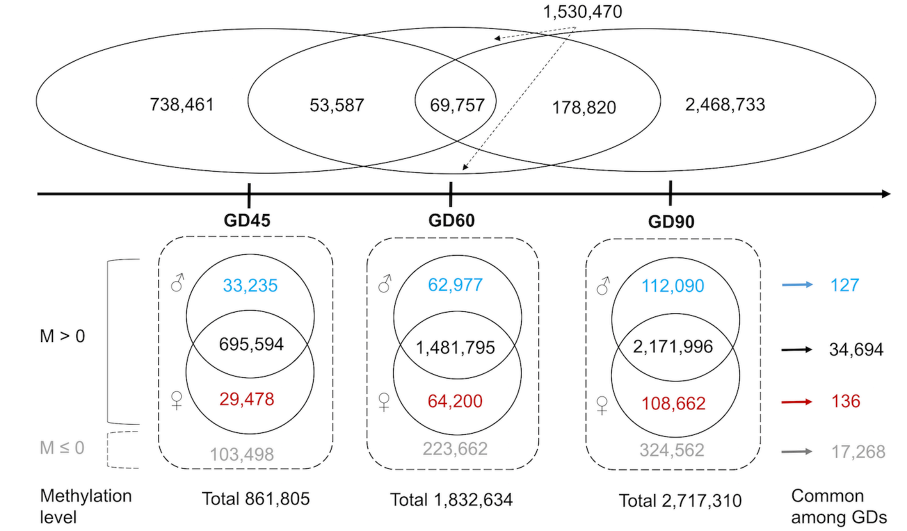

### Supplementary Figure 2

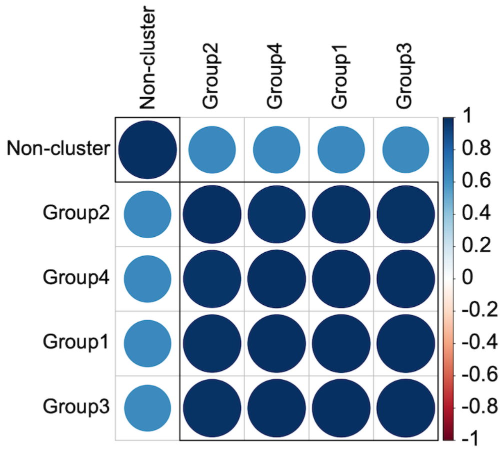

### Supplementary Figure 3

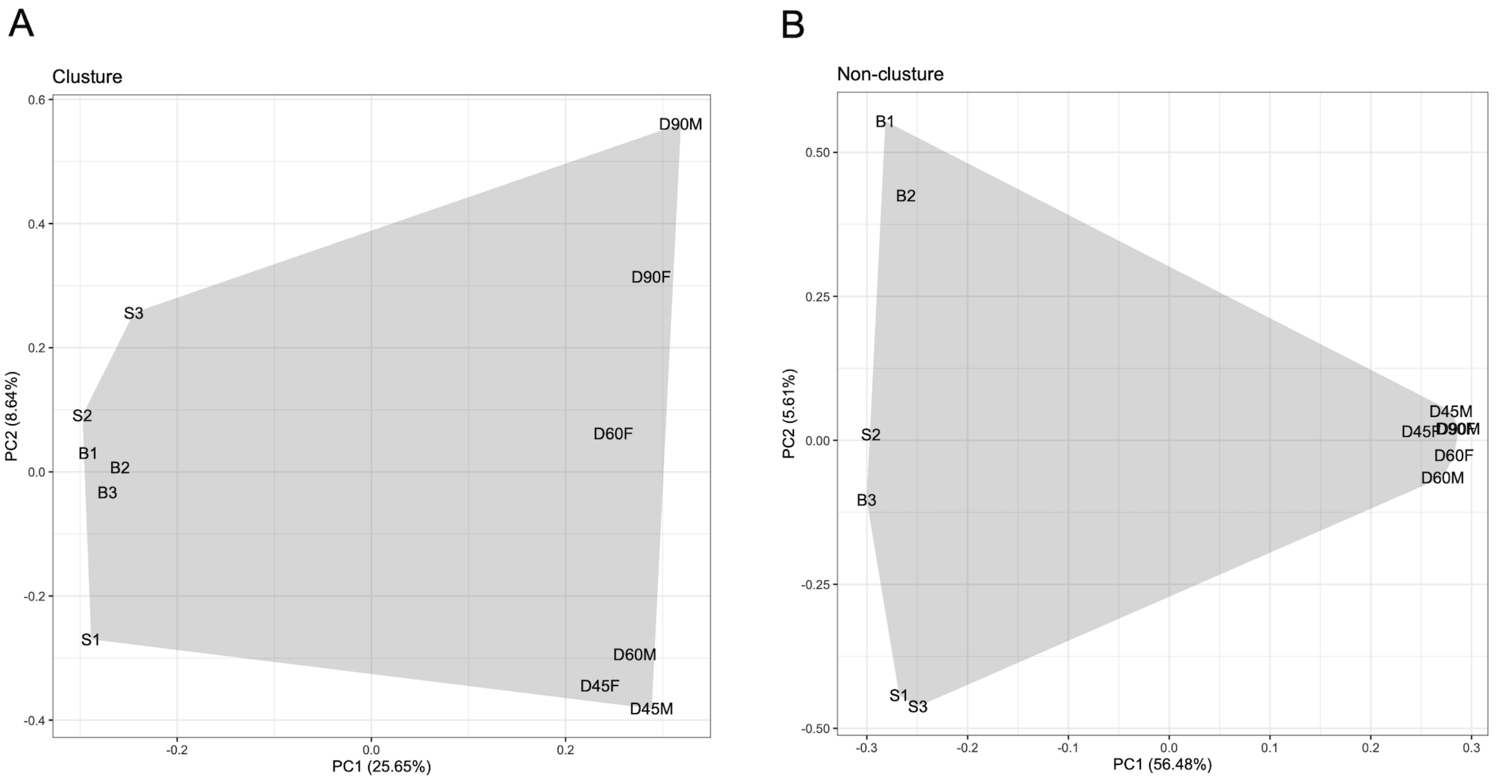

### Supplementary Figure 4

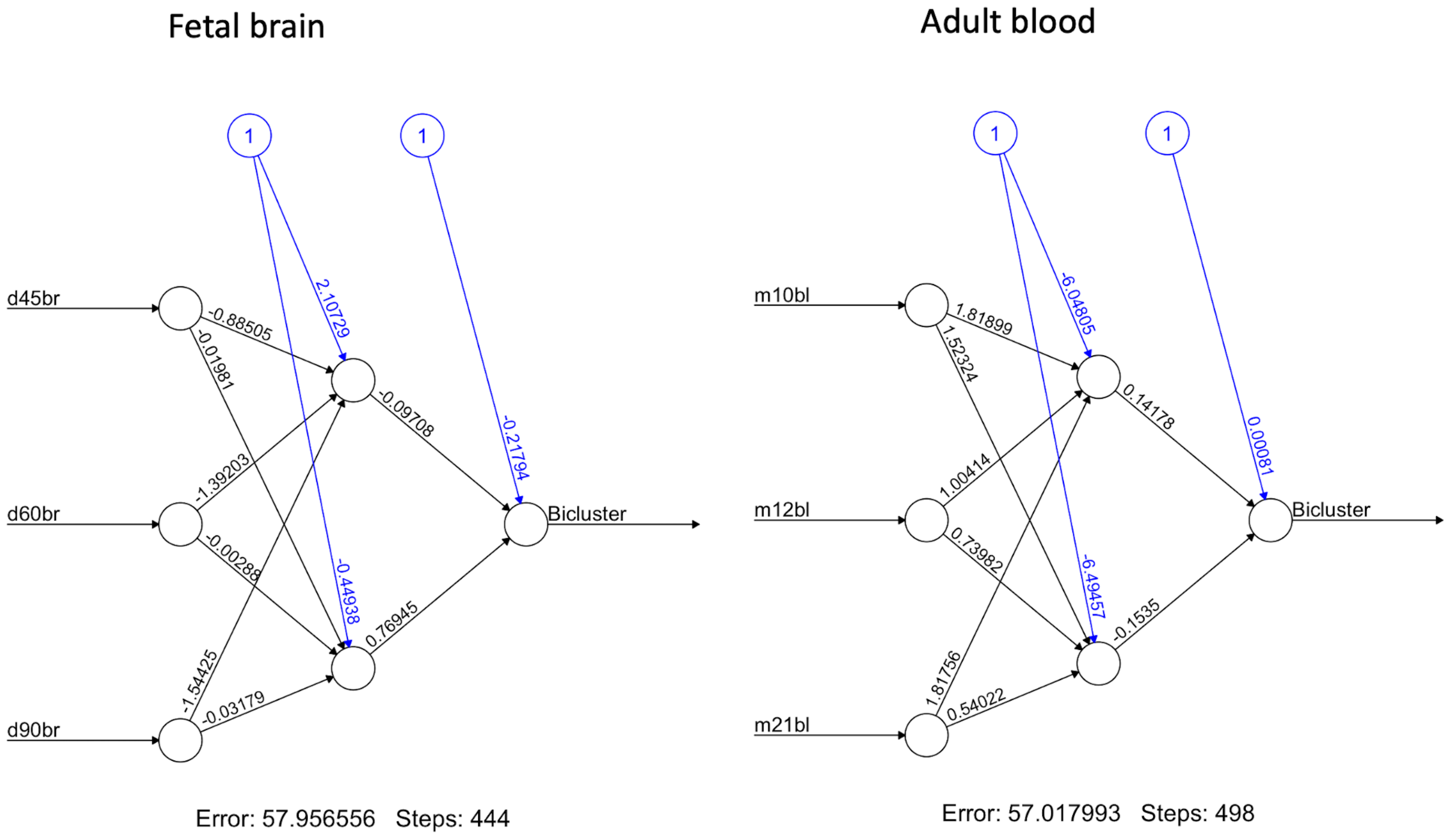

### Supplementary Figure 5

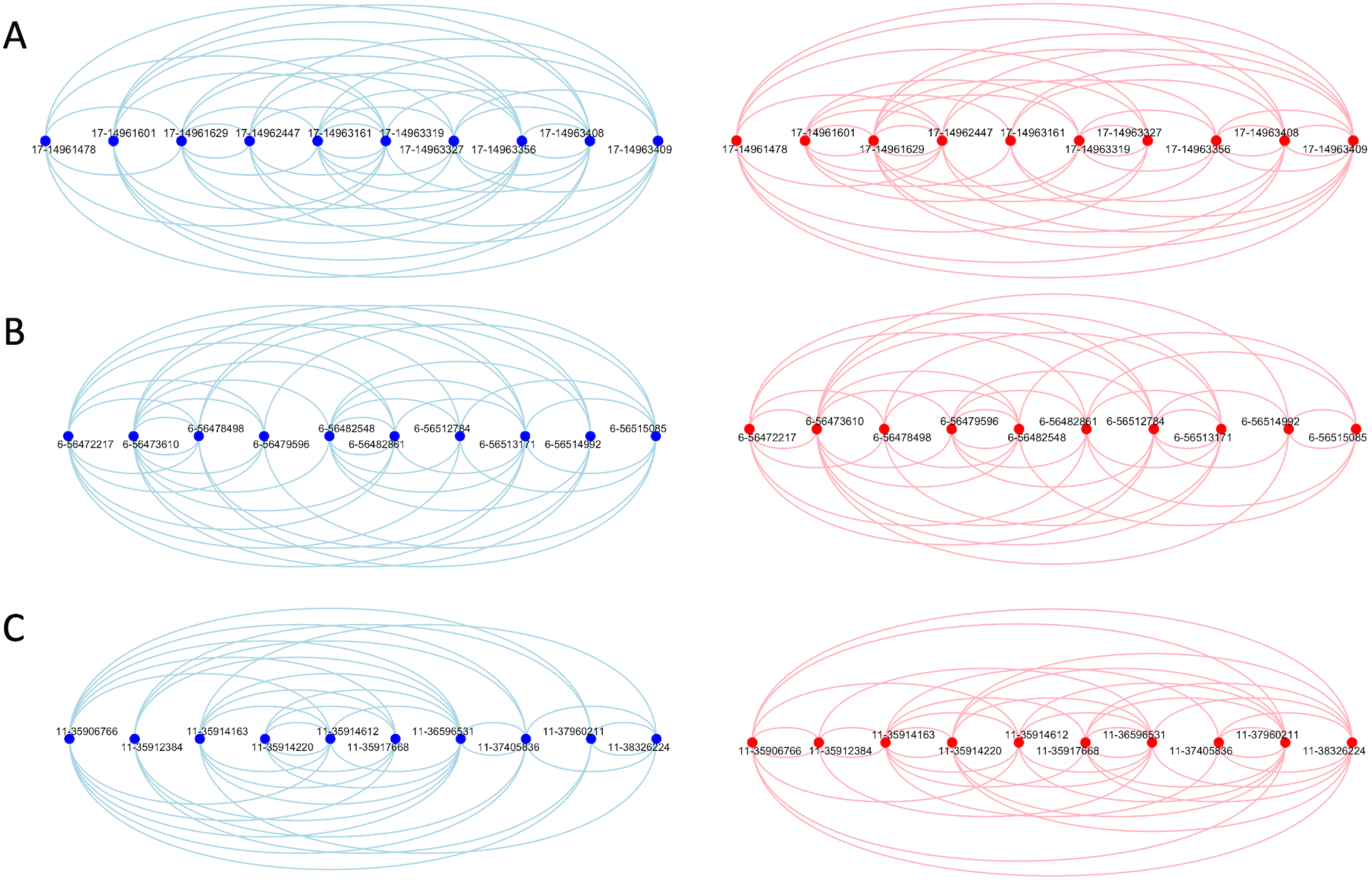


Fetal brain

Adult blood
